# MarcoPolo: a clustering-free approach to the exploration of differentially expressed genes along with group information in single-cell RNA-seq data

**DOI:** 10.1101/2020.11.23.393900

**Authors:** Chanwoo Kim, Hanbin Lee, Juhee Jeong, Keehoon Jung, Buhm Han

## Abstract

A common approach to analyzing single-cell RNA-sequencing data is to cluster cells first and then identify differentially expressed genes based on the clustering result. However, clustering has an innate uncertainty and can be imperfect, undermining the reliability of differential expression analysis results. To overcome this challenge, we present MarcoPolo, a clustering-free approach to exploring differentially expressed genes. To find informative genes without clustering, MarcoPolo exploits the bimodality of gene expression to learn the group information of the cells with respect to the expression level directly from given data. Using simulations and real data analyses, we showed that our method puts biologically informative genes at high ranks more robustly than other existing methods. As our method provides information on how cells can be grouped for each gene, it can help identify cell types that are not separated well in the standard clustering process. Our method can also be used as a feature selection method to improve the robustness of the dimension reduction against changes in the parameters involved in the process.

## Introduction

Single-cell RNA (scRNA) sequencing technology has offered opportunities to study gene expressions of individual cells of a biological system. A common approach to analyzing the data in the first place is to perform an unsupervised clustering method to group cells^1,2^. Then based on the clustering result, differentially expressed genes (DEGs) between the groups of cells are identified^3,4^. Since people typically consider only the top-ranked DEGs in the analysis, in case the grouping is wrong, it is nearly impossible to rediscover other missed but informative genes^5^. Thus, the interpretation of data via downstream differential expression analysis largely depends on the predetermined clustering result.

However, a clustering result has an innate uncertainty. Since clustering depends on a specific algorithm, it is unknown whether the clustering result appropriately reflects the underlying biological structure of the data^6,7^. For this reason, it is common that the clustering procedure is performed repeatedly to get the seemingly best clustering result^3^. At each trial, one performs clustering with a single parameter setting and checks the soundness of clustering by comparing cluster-level expression profiles with the list of already known marker genes. These two steps can be repeated with varying parameters until the identity of each group is confidently determined. However, there are numerous options and parameters, such as the methods for highly variable genes (HVGs) selection^3,4,8^, the number of HVGs used^3,8^, the methods for dimensionality reduction^9,10^, and the parameter for the clustering resolution^2,11^. Thus, the process is arduous in many cases and picking the best observation from a large number of trials may often lead to data-specific overfitting. Furthermore, our prior knowledge of marker genes is incomplete, making it hard to conclude that the obtained clustering result represents the ground truth^7,12^.

In this respect, there have been demands for methods that can extract differentially expressed genes from data without being susceptible to uncertainty rooting from clustering results^5,13^. One possible approach is to utilize existing HVG methods. However, HVG methods were designed to select genes as input for the dimensionality reduction step, but were not designed to sort out DEGs. In our analysis, we show that they often do not assign high ranks to DEGs that are obviously differentially expressed among cell types with sufficient precision compared to the standard DEG analysis with clustering. Another advanced approach is singleCellHaystack, a recently developed method that extracts a list of candidates for DEGs by examining non-random expression pattern^14^. It examines whether the expression of a gene agrees with the placement of cells in low-dimensional space such as PCs (i.e., local cell density). Although this method sorts out DEGs in a more sophisticated way than the HVG methods, similarly to HVG methods, it does not tell which groups of cells ‘differentially’ express the identified genes. Accordingly, the process of manually examining the expression pattern of genes is left to users. In addition, as the authors pointed out as a weak point, the method requires the use of a predetermined threshold to obtain binary detection data (i.e., whether a gene is either expressed or not in each cell); However, the use of hard thresholding can lead to considering two subsets of cells as a single group even if both express a gene but with different expression intensities. It is common that expression between two subsets of cells may not be binary but bimodal^15–17^.

We here propose MarcoPolo, a novel clustering-free approach to identifying DEGs in scRNA-seq data. The two main functions of our method are to sort out genes with biologically informative expression patterns with high precision without clustering, and to learn tentative grouping of the cells with respect to the expression level (which groups of cells ‘differentially’ express the identified genes) directly from the given data. As our method does not demand prior clustering information of cells in advance, our approach is robust to uncertainties from clustering or cell type assignment. Additionally, our framework provides the analysis result in the form of an HTML file so that researchers can conveniently interpret and make use of the result for various purposes.

MarcoPolo achieves a high precision in finding informative genes by using the following strategies. It first disentangles the bimodality inherent in gene expression and divides cells into two groups by the maximum likelihood estimation under a mixture model. Thus, it takes advantage of the fact that the difference of expression patterns of a gene between two subsets of cells can be bimodal.^15–17^ Then, it goes through additional processes to confirm which genes have a differential expression pattern that is biologically feasible. Specifically, it uses a *voting system* that compares how cells are grouped for a gene with how cells are grouped for other genes in order to see if the groupings are similar. This is based on a biological phenomenon that a group of cells in a similar biological status co-expresses a subset of genes dependently altogether^18^. Hence, if the grouping pattern of a gene is repeated, or supported, by many other genes, the gene is considered informative. In addition to this strategy, MarcoPolo uses a statistic examining whether the cells expressing a gene are proximal in the reduced dimension space. MarcoPolo combines these three scores (bimodality score, voting system, and proximity score) into one to winnow genes of which expressions are noteworthy. Using extensive simulations and real data analyses, we demonstrate that MarcoPolo can have utilities for identification of informative genes, detection of interesting groups of cells, and selection of feature genes for downstream clustering analysis.

## Results

### Overview of MarcoPolo Method

MarcoPolo is a mixture-model-based multiple-criteria ranking approach to selecting differentially expressed genes. MarcoPolo fits a two-component Poisson mixture model and ranks the genes using the parameters estimated from the fitted model (**Methods**). For a given marker, cells that are more likely to be part of the high expression component of the mixture distribution are named on-cells while those that are more likely to be part of the low expression component are named off-cells (**Figure 1A**). We developed a ranking method called the *voting system* that prioritizes genes that exhibit a common expression pattern with other genes (**Figure 1B**). The intuition behind this ranking method is that a true biological entity will express multiple markers; therefore, a gene that reflects this true entity will have other genes with a similar expression pattern. For example, gene 1, gene 3, gene 4, and gene 5 in **Figure 1B** show similar on-cell patterns. Therefore, the voting system assigns higher ranks to these genes. By contrast, gene 2 and gene 6 have no other genes sharing on-cell patterns similar to theirs and thus are assigned lower ranks by the voting system. In addition to the voting system, we implemented two more complementary scores, the *proximity score* and the *bimodality score.* We expect that cells within the same cluster will be close in the distance in the low-dimensional representation space. Based on this idea, the proximity score computes the variance of the principal component (PC) values of on-cells and assigns higher ranks to genes with low variance (**Figure 1C**). The bimodality score system measures how well the bimodal two-component model fits to a given gene (**Figure 1D**). This system examines two things. First, this system compares the log likelihood of the model (Q-score) of the two-component model versus the one-component model. Accordingly, genes with a bigger reduction in the Q-score are ranked higher by the bimodality score ranking system. In addition, the bimodality score ranking system assigns higher ranks to genes of which on-cells’ mean expression value is much higher than all cells’ mean expression value. Finally, MarcoPolo determines the ranking of all genes by combining these three scoring systems. Our framework also provides the analysis result in the form of an HTML file so that researchers can conveniently interpret and make use of the result for various purposes (**Figure 1E**). For each gene, the output file provides fold change based on the tentative group, a two-dimensional plot of cells, a histogram of expression with annotated group information, and the statistics that were used to winnow genes.

**Figure 1.**
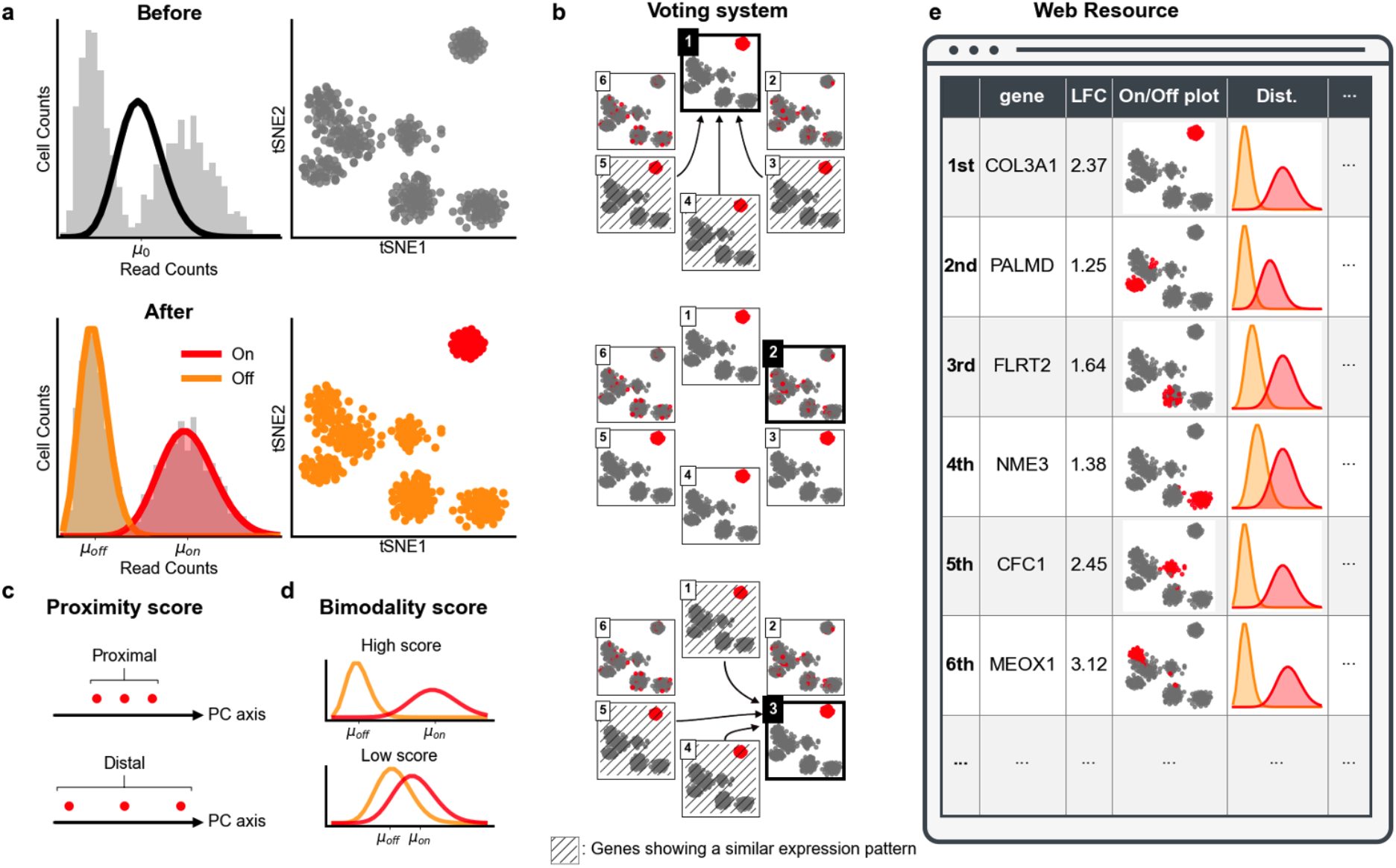
Overview of MarcoPolo. **(a)** MarcoPolo fits a two-component Poisson mixture model. In t-SNE plot, on-cells, which are more likely to be part of the high expression component of the mixture distribution, are colored red. Off-cells, which are more likely to be part of the low expression component of the mixture distribution, are colored orange. **(b)** In voting system, genes exhibiting a common expression pattern with other genes are prioritized. For each gene, the on-off assignment of the gene is compared with those of other genes. Genes that are supporting the compared one are indicated using arrow. Thus, gene 1 and gene 3 are supported three times. In contrast, gene 2 in the middle is not supported by any of genes shown. **(c)** In proximity score system, the variance of the principal component (PC) values of on-cells is calculated for each gene. Higher ranks are assigned to genes with low variance. **(d)** In bimodality score system, genes with a bigger reduction in the Q-score or of which on-cells’ mean expression value is much higher than all cells’ mean expression value are ranked higher. **(e)** MarcoPolo framework offers the analysis result in the form of an HTML report. For each gene, log fold change, On/Off plot, biological description, and statistics calculated by MarcoPolo are shown. On/Off plot shows how cells can be grouped for each gene according to the bimodality in its expression.

### Clustering uncertainty can hinder the discovery of differentially expressed genes

As the differential expression analysis in downstream depends on the clustering result, if a clustering algorithm fails to properly cluster a group of cells, the informative genes of that cluster will be missed. In this sense, the need for a method that can find DEGs independent of clustering was widely discussed^5,13,14^. We demonstrated this situation as a proof-of-concept using realistic simulation datasets generated by Symsim^19^, a simulator of single-cell RNA-seq experiments. Varying the probability that a gene has a nonzero type-specific expression effect size in the simulator, we generated 40 different simulation datasets (**Methods**). As expected, the more the type-specific gene effect sizes were turned off, the less clear the boundaries between the cell populations became (**Figure 2A**), and the lower the quality of clustering results became (**Supplementary Figure 1**). For the generated datasets, we compared MarcoPolo with the standard DEG workflow with clustering (performing Seurat clustering with default parameters and finding markers with findMarker function of Seurat), two widely used HVG methods implemented in Seurat package^11^ (VST and DISP), and singleCellHaystack. We used the area under the receiver operating characteristic (ROC) curve to see how accurately each method sort out marker genes (**Figure 2B**), for which non-zero true effect sizes were assumed. We found that the performance of the standard workflow with clustering was negatively affected by the decrease of this parameter (non-zero effect gene proportion) as expected. When the probability of non-zero type-specific expression effect size was set as 1e-2, the medians of standard workflow’s AUCs were relatively high (0.934~0.978), and the inter-quartile range (IQRs) of them were small (0.048~0.066). However, when the parameter was lowered to 5e-4, the medians went down (0.764~0.847), and the IQRs became large (0.154~0.309). Interestingly, the performance of singleCellHaystack showed a similar trend to the standard workflow. When the parameter was 1e-2, the median was relatively high (0.942), and the IQR was small (0.032); but when the parameter was 5e-4, the median went down (0.788), and the IQR became large (0.378). In contrast, MarcoPolo and HVG methods were more robust to these changes. Their medians did not go below 0.835, and the IQRs remained all below 0.110. Although the simulation datasets do not perfectly represent real situations, this result suggests that when it is hard to cluster cells appropriately, the standard DEG analysis with clustering could easily miss the target populations’ DEGs that could be retrieved by MarcoPolo or simply using HVG methods.

**Figure 2.**
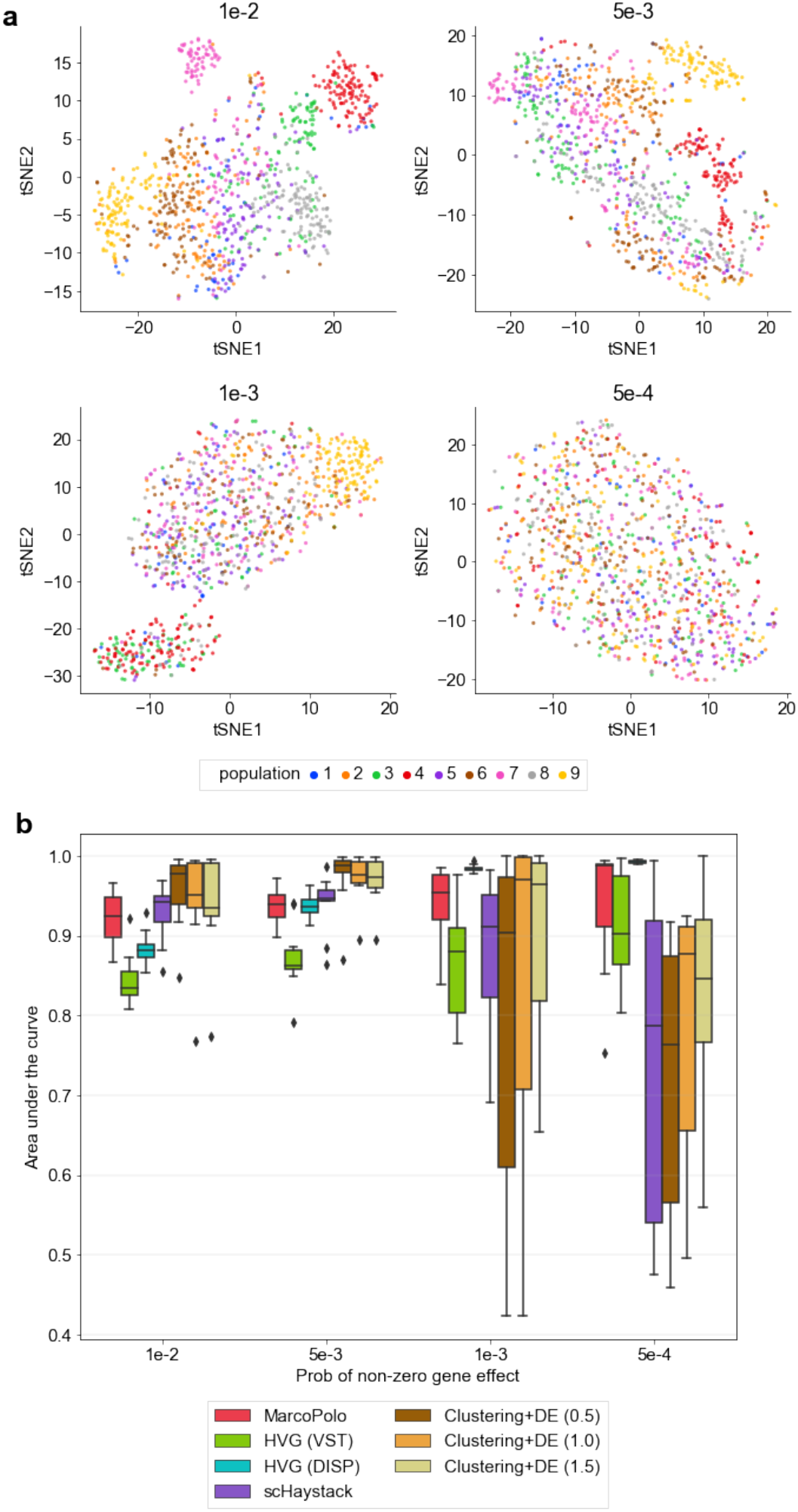
Comparison of methods applied to simulation datasets. **(a)** t-SNE plots of simulation datasets when changing the probability of non-zero type-specific expression effect sizes. **(b)** the area under the curve of the gene lists generated by different methods when changing the probability of non-zero typespecific expression effect sizes. The gene lists were obtained by using MarcoPolo, HVG methods, singleCellHaystack, or standard DEG pipeline with clustering. For standard DEG pipeline, the number inside parenthesis denotes the resolution parameter of the clustering algorithm.

### MarcoPolo puts genes with biologically feasible expression patterns at the top

We applied MarcoPolo to 23 real datasets. We used the human embryogenic stem cell (hESC) dataset^20^, human liver cell dataset^21^, and human peripheral blood mononuclear cell (PBMC) dataset^22^, and Tabula Muris consortium datasets of 20 different organs and tissues^23^. In these datasets, the cells were processed either by fluorescence-activated cell sorting (FACS) or by manual curation based on known markers. In our analysis, we assumed that the cell-type labels provided by their authors are correct and defined genes that are expressed cell-type-specifically, namely the cell-type markers, based on these cell type labels. Then, we checked how well different methods put the true markers at the top ranks if the scRNA-seq data was given without the true type labels (**Methods**). Although the method showing the best performance differed by datasets, overall, MarcoPolo showed comparable or better performance in sorting out marker genes than the standard DEG analysis with clustering (**Figure 3** and **Supplementary Figure 2**). The median AUC was 0.850 for MarcoPolo, 0.772 for the DISP (HVG), 0.665 for VST (HVG), 0.772 for singleCellHaystack, and around 0.770~0.774 for standard DEG pipelines with clustering. In addition, compared with other methods, the inter-quartile range of MarcoPolo was relatively small (0.128). On top of that, when we looked at which method performed the best for each dataset, MarcoPolo performed the best or the second-best in 19 out of 23 real datasets (83% of datasets; 12 best and 7 second-best, respectively) (**Supplementary Figure 2**). For singleCellHaystack, the number was 8 (35% of datasets; 5 best and 3 second-best respectively). For standard pipeline with clustering, the number 6 (26% of datasets; 2 best and 4 second-best respectively). This result shows that MarcoPolo’s multiple-criteria ranking approach works well in sorting out informative genes for a variety of datasets. As MarcoPolo determines the final ranking of the genes by combining the three scoring systems, we wanted to see how each score system in MarcoPolo contributed to the final performance. To this end, we examined the performance of using each score system alone (**Supplementary Table 1**). The scoring system with the best performance differed by datasets, showing that MarcoPolo needed all three systems to achieve good performance over different datasets. The performance of the combined final score was often similar the best performing single system, which demonstrated that the combining strategy was effectively capturing the information from the best system for each dataset.

**Figure 3.**
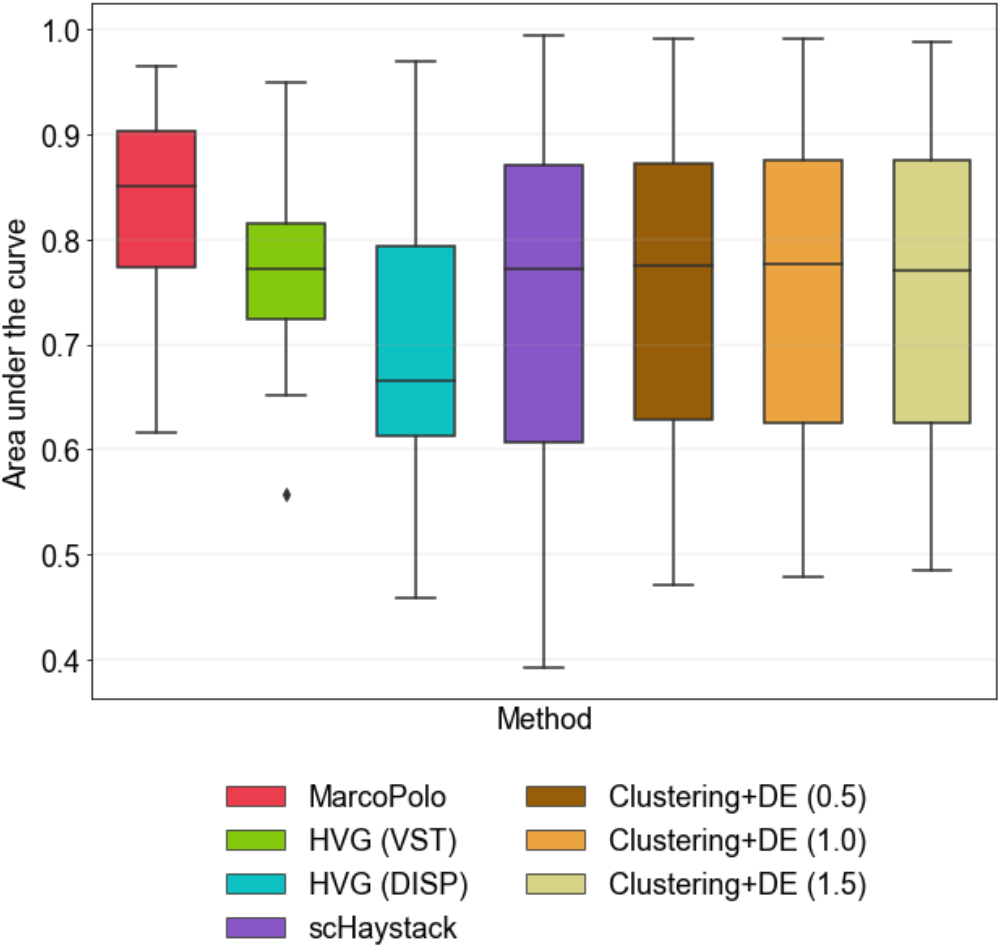
Comparison of methods applied to real datasets. The area under the curve of the gene lists is shown. The gene lists were obtained by using MarcoPolo, HVG methods, singleCellHaystack, or standard DEG pipeline with clustering. For standard DEG pipeline, the number inside parenthesis denotes the resolution parameter of the clustering algorithm.

### Genes found by MarcoPolo can suggest cell types via bimodal expression patterns

Compared with singleCellHaystack, which uses a fixed threshold to determine whether a gene is either expressed or not in each cell, MarcoPolo adapts flexible thresholds to each gene’s expression pattern. Thus, even if two subsets of cells express a gene at the same time, in case they have bimodal expression patterns, MarcoPolo can classify them into two separate groups. For a given marker, cells that are more likely to be part of the high expression component of the mixture distribution are grouped as on-cells while those that are more likely to be part of the low expression component grouped as off-cells. As an example, how cells in datasets were divided by MarcoPolo was shown (**Figure 4A-C**). For the genes shown, a certain cell type showed a higher level of expression intensity than other cell types. MarcoPolo successfully identified them as a separate group. If MarcoPolo had used a fixed threshold, multiple cell types might have been identified as a single group.

**Figure 4.**
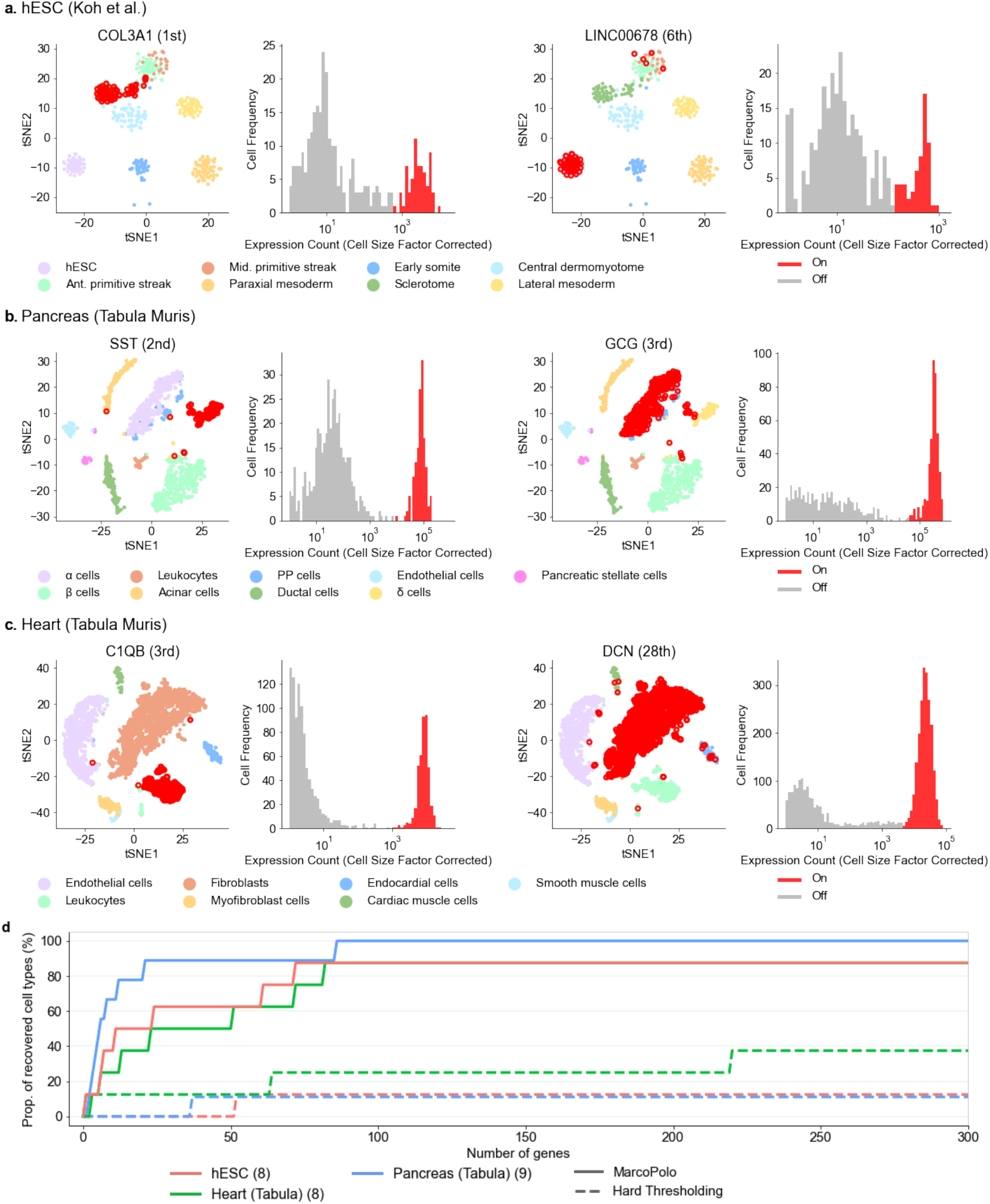
The identification of cell types by using MarcoPolo’s grouping information. **(a)-(c)** how cells in each dataset were divided by MarcoPolo. For a given marker, cells that are more likely to be part of the high expression component of the mixture distribution are grouped as on-cells while those that are more likely to be part of the low expression component grouped as off-cells. t-SNE plots of each dataset were labeled by the true cell-type labels. The number beside the name of each gene is its rank in MarcoPolo result. Each histogram shows the expression pattern of a gene in the dataset. Cells with zero expression count were excluded from plotting. The edge of dots that corresponds to on-cells for each gene was colored red. The expression count was divided by cell-specific scaling factors (also known as size factors). **(d)** The proportion of recovered cell types in the datasets with the increasing number of genes reviewed in MarcoPolo result. The grouping information of each gene was reviewed according to its rank assigned by MarcoPolo. The number inside the parenthesis beside the dataset name denotes the number of cell types in each dataset.

As MarcoPolo provides information on how cells can be grouped for each gene, one can use MarcoPolo to identify cell types in the given data without the help of a clustering algorithm. It is true that not all cell types have a marker gene of which bimodal expression is significant to identify them and that accordingly, for some cell types, signals from multiple genes need to be considered. However, we found that MarcoPolo was able to retrieve a substantial amount of cell types in the real datasets by exploiting the bimodal expression patterns of genes (**Figure 4D** and **Supplementary Figure 3**). We measured how many cell types can be retrieved by MarcoPolo’s grouping information and how many genes at the top ranks in MarcoPolo should be reviewed to achieve this (**Methods**). We regarded a gene to be distinguishing a cell type if its on-cells specifically indicated the cell type (i.e., if the on-cell group contains more than 70% of the cell type and less than 20% of the other remaining cell types). In total, among the 147 cell types included in 23 real datasets, MarcoPolo’s grouping information was able to segregate 97 cell types when the top 300 genes in MarcoPolo were considered (**Supplementary Table 2**). Obviously, the grouping by hard thresholding (i.e., binary detection data) failed to identify cell types when the same procedure was followed.

### MarcoPolo genes can distinguish cell types that are not distinguished by the standard pipeline

MarcoPolo succeeded in identifying cell types that were not distinguished by the standard pipeline. The human embryogenic stem cell (hESC) dataset of Koh et al.^20^, the lung dataset of Tabula Muris consortium^23^, and the liver dataset of MacParland et al.^21^ are examples showing that the standard clustering approach can often fail to distinguish true cell types. In the precalculated 2D t-SNE coordinates that were included in the downloaded datasets, the boundaries between the anterior primitive streak (APS) and mid primitive streak (MPS) in the hESC dataset, natural killer (NK) cells and T cells in the lung dataset, and gamma delta T cells and NK cells in the liver dataset were unclear (**Supplementary Figure 4**). Since t-SNE reflects the variance in the PC space, this mixture suggests that PC space-based clustering algorithms will likely fail to distinguish them as well.

We tried running a clustering algorithm on the datasets from scratch using the standard Seurat pipeline (**Figure 5**). We used the default setting (VST method to select 2,000 genes) to calculate PCs. For the clustering step, we used the widely used option (FindNeighbors and FindClusters function in Seurat package). For the FindClusters function’s resolution parameter, we used the value of 2.0 (default: 1.0) to get a fine clustering result. In our analysis, similarly to the precalculated t-SNE coordinates, the clustering algorithm failed to group the abovementioned cell types in each dataset properly. This implies that, if one simply uses the standard pipeline, one would have difficulties in distinguishing these cell types and finding DEGs between them.

**Figure 5.**
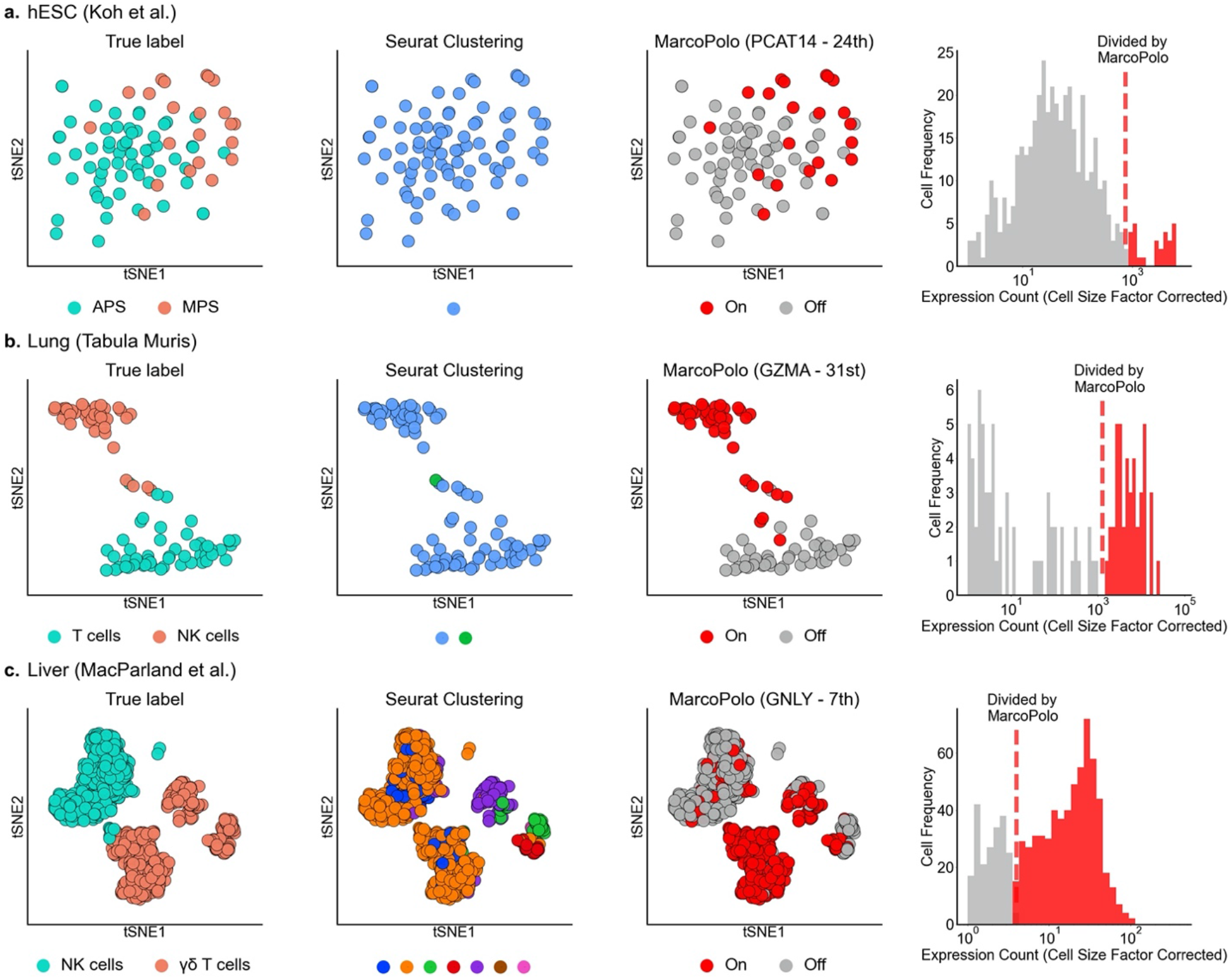
Cell types distinguished by MarcoPolo genes. They are cell types that were not distinguished using the default option of standard clustering pipeline. Exceptionally, we used the clustering resolution parameter of 2.0 (default: 1.0) to obtain a finer clustering result. We plotted cells on the precalculated t-SNE coordinates that were included in the downloaded datasets. For each dataset, we only showed cell types of interest. In the t-SNE plot in the first column, we colored cells by their true cell-type label. In the second t-SNE plot, we colored cells by Seurat clustering result. In the third t-SNE plot, we colored cells based on MarcoPolo grouping. For each gene, MarcoPolo learns how to divide cells in the given dataset into two groups according to their expression modality. The cells belonging to the on-cell group (i.e., the group of higher gene expression) were colored red. In the histogram, the expression pattern of each gene was shown. Cells with zero expression count were excluded from plotting. The expression count was divided by cell-specific scaling factors (also known as size factors). **(a)** the hESC dataset of Koh et al. **(b)** the lung dataset of Tabula Muris consortium **(c)** the liver dataset of MacParland et al.

We extracted DEGs based on the true cell-type labels (i.e., the APS and MPS for the hESC dataset) and checked how well each method identifies them (**Table 1**). For each dataset, we calculated fold change between each unseparated cell type and all other cell types and chose the top 3 DEGs regardless of two cell types in the order of fold change. Compared with singleCellHaystack and standard DEG pipeline with clustering, MarcoPolo and HVG methods put those DEGs at the top ranks well. In addition to pointing out the genes well, MarcoPolo provided grouping identifying each cell type highly accurately (**Figure 5** and **Supplementary Figure 5**). For example, in the hESC dataset, MarcoPolo successfully identified MPS and APS as two different groups based on the expression of the PCAT14 gene, which ranked at the top 24th.

**Table 1.** Differentially expressed genes based on true cell-type labels. Three datasets contained cell types that are not distinguished using the default option of standard clustering pipeline. They are the anterior primitive streak (APS) and mid primitive streak (MPS) in the hESC dataset, natural killer (NK) cells and T cells in the lung dataset, and gamma delta T cells and NK cells in the liver dataset, respectively. For each dataset, we calculated fold change between each unseparated cell type and all other cell types and chose the top 3 DEGs regardless of two cell types in the order of fold change. We examined how each gene was ranked by different methods.

| FC-rank | LFC | Gene | Cell type | MarcoPolo | HVG (VST) | HVG (DISP) | scHaystack | C+DE (0.5) | C+DE (1.0) | C+DE (1.5) |
| --- | --- | --- | --- | --- | --- | --- | --- | --- | --- | --- |
| **hESC (Koh et al.)** |  |  |  |  |  |  |  |  |  |  |
| 1st | 2.66 | PCAT14 | MPS | 24th | 3rd | 29th | 6957th | 1427th | 1427th | 1427th |
| 2nd | 2.07 | CER1 | APS | 72nd | 7th | 51st | 4th | 36th | 36th | 36th |
| 3rd | 1.66 | NODAL | APS | 9th | 766th | 621st | 46th | 5th | 5th | 5th |
| **Lung (Tabula Muris)** |  |  |  |  |  |  |  |  |  |  |
| 1st | 5.03 | GZMA | NK cells | 31st | 1st | 12nd | 1511st | 552nd | 481st | 442nd |
| 2nd | 4.20 | CCL5 | NK cells | 41st | 2nd | 26th | 1613rd | 501st | 434th | 398th |
| 3rd | 4.09 | NKG7 | NK cells | 39th | 8th | 39th | 1254th | 195th | 127th | 125th |
| **Liver (MacParland et al.)** |  |  |  |  |  |  |  |  |  |  |
| 1st | 3.50 | GNLY | $\gamma\delta$ T cells | 7th | 45th | 42nd | 625th | 28th | 22nd | 24th |
| 2nd | 2.48 | CMC1 | NK cells | 64th | 253rd | 95th | 372nd | 64th | 62nd | 374th |
| 3rd | 2.36 | GZMB | $\gamma\delta$ T cells | 153rd | 116th | 167th | 1035th | 124th | 112nd | 108th |

Although two of the cell types all expressed the gene, using the differences in their expression intensity, MarcoPolo was able to distinguish them.

### Using MarcoPolo as a feature selection method to improve the robustness of the clustering process

In the standard scRNA-seq analysis pipeline, a subset of genes selected by HVG methods is used to construct a low dimensional representation for the clustering step. To obtain a clustering result that well reflects the structure of the underlying biological data structure, it is important to use informative genes as an input. As MarcoPolo and singleCellHaystack are designed to pick genes with informative differential expression, we used them as a feature selection method in the standard pipeline. We mixed MarcoPolo (or singleCellHaystack) genes with HVGs and used them as input for the dimensionality reduction step. We wanted to see if the robustness of the clustering procedure improved when this approach is taken. We defined the robustness of the clustering procedure as the frequency of successfully clustering cells correctly with varying parameters and settings. For each clustering result, we observed whether cell types discussed in the previous section (the APS and MPS in the hESC dataset, NK cells and T cells in the lung dataset, and γö T cells and NK cells in the liver dataset) were successfully distinguished.

Briefly, we compared three categories of feature selection methods: only using genes selected by standard HVG method, using the mixture of HVG genes and MarcoPolo genes (namely HVG with MarcoPolo), and lastly using the mixture of HVG genes and singleCellHaystack genes (namely HVG with Haystack). In case we mixed two different criteria such as HVG and MarcoPolo, we extracted the same number of genes from the top-ranked genes in each criterion.

In order to test the robustness of each feature selection method, we ran the same method multiple times using different parameters and settings. Then, we measured how many times a method gave a successful clustering over the wide range of tested parameters. Specifically, we varied the parameters and settings as follows. First, the HVG method was used in all three methods (that is, it was used as a standalone or in a mixture). Although there are different HVG methods available, we tried two widely used methods (VST and DISP) implemented in the Seurat package. Second, we varied the number of HVGs from 200 to 1,000 with the interval of 100 (9 numbers). Third, we varied the number of top PCs used by the clustering algorithm from 10 to 30 with the interval of 5 (5 numbers). Fourth, we varied the resolution parameter in the Louvain clustering algorithm from 0.6 to 1.4 with the interval of 0.2 (5 parameters). To sum up, for each category of feature selection methods, we repeatedly clustered cells 450 times with different settings (2 HVG methods × 9 HVG numbers × 5 top PC numbers x 5 resolution parameters). We then calculated how many of those 450 trials succeeded in isolating the populations.

Overall, employing MarcoPolo as a feature selection method improved the frequency of successful clustering (**Figure 6A**). When MarcoPolo genes were employed for the hESC dataset, the frequency of separating the populations improved from 36.4% to 85.6% compared to using HVGs alone. For the lung dataset, it improved from 41.8% to 56.9%. For the liver dataset, it slightly lowered from 78.7% to 76.2%. singleCellHaystack genes also showed similar trends but had a larger performance drop in the liver dataset. When it comes to the overall performance of all the datasets, HVG with MarcoPolo, HVG with singleCellHaystack, and HVGs were 72.9%, 60.0%, and 52.3%, respectively. We also compared the methods separately for each parameter of the clustering pipeline (**Figure 6B-D**). MarcoPolo performed the best in many cases. In addition, in the figures, the performance of methods was shown separately for different datasets. We observed that, in general, MarcoPolo was the best choice to increase the probability of separating the cell types.

**Figure 6.**
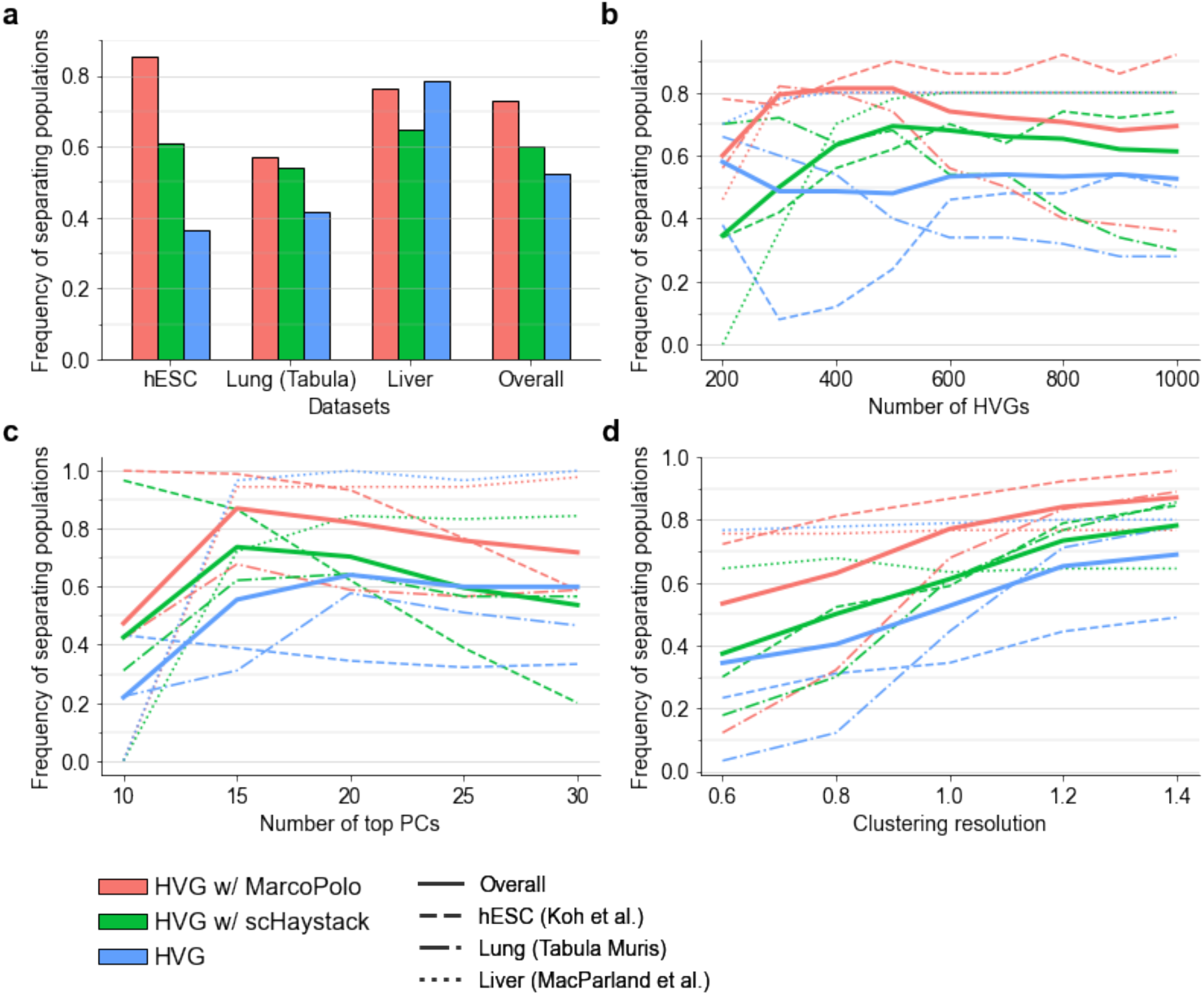
Frequency of successfully separating populations of interest over multiple trials with varying parameters. Using each feature selection method, we selected genes that are used by the downstream clustering analysis. We tried 450 different settings and parameters for the clustering analysis and measured the frequency of successful clustering. **(a)** Comparing the performances according to the different datasets. **(b)-(d)** Comparing the performances according to each parameter of the clustering pipeline. **(b)** Comparing the performances according to varying the number of HVGs. **(c)** Comparing the performances according to varying the number of top PCs. **(d)** Comparing the performances according to varying resolution parameter of the clustering algorithm.

## Discussion

We showed that the MarcoPolo approach has three practical usages. It can identify biologically informative genes accurately. It can detect interesting groups of cells that are not identified well in the standard clustering process. Additionally, it can be used as a feature selection method for downstream clustering analysis.

Compared with the standard DEG analysis with clustering, MarcoPolo is totally new approach to analyzing scRNA-seq data. In the standard workflow, groups of cells are defined first, and then DEGs among the groups are identified. In contrast, MarcoPolo first identifies DEGs and then provides tentative grouping information for each gene. It is true that not all cell types have a marker gene of which bimodal expression is significant to identify them and that accordingly, for some cell types, signals from multiple genes need to be considered. However, in our analysis, MarcoPolo’s grouping information was shown to recover a substantial number of cell types in real datasets. It was even able to distinguish cell types that were not distinguished by the standard pipeline.

In addition to providing grouping identifying each cell type highly accurately, MarcoPolo put genes with noteworthy gene expression patterns at the top. It first disentangles the bimodality inherent in gene expression and divides cells into two groups by the maximum likelihood estimation under a mixture model. Then, it measures three kinds of statistics to confirm which genes have a differential expression pattern that is biologically feasible. We showed that the multiple-criteria ranking approach worked well in sorting out informative genes for a variety of datasets.

In relation to other methods, the contribution of our method is as follows. Our contribution is that by giving an appropriate amount of flexibility to defining cell clusters, MarcoPolo can lower the risk of missing informative genes. Our analysis compared four kinds of approaches to discovering DEGs: HVGs, singleCellHaystack, MarcoPolo, and the standard workflow with clustering. Compared with HVGs, singleCellHaystack takes into account other genes’ expression to determine each gene expression’s nonrandomness. It uses low-dimensional data to confirm that each gene’s expression agrees well with the given dataset’s overall variance. MarcoPolo is located somewhere in between singleCellHaystack and the standard workflow with clustering. To determine each gene expression’s biological feasibility, similarly to singleCellHaystack, MarcoPolo also considers other genes’ expression in the given data but in a more sophisticated way than singleCellHaystack. It estimates tentative grouping information for each gene. Then, it determines whether each gene’s expression well agrees with the given data’s overall expression pattern with the tailored grouping information for the gene. In this way, MarcoPolo discovers DEGs more accurately. Compared with the standard workflow that uses fixed membership of cells, MarcoPolo has more flexibility in defining cell clusters as it uses tentative grouping per each gene. Using this approach, MarcoPolo distinguishes cell types that are not distinguished by the standard pipeline.

## Conclusion

We presented MarcoPolo, a clustering-free approach to the exploration of bimodally expressed genes in single-cell RNA-seq data. Our method exploits the bimodality of gene expression to learn group information from the data while finding informative genes without clustering. Using simulations and real data analyses, we showed that our method has advantages as follows. First, our method puts genes with biologically informative expression patterns at the top ranks accurately and robustly. Second, as our method provides information on how cells can be grouped for each gene, it can help identify cell types that are not separated well in the standard clustering result. Third, our method can also be used as a feature selection method to improve the robustness of the clustering process.

## Methods

### Linear Poisson mixture model

To identify the expression modality in scRNA-seq data, we fit the following Poisson mixture model to each gene’s count data.

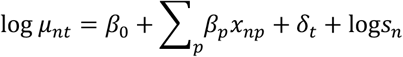

Here, *t* ∈ {0,1} indicates the two groups, the on-cells and off-cells. Conditional on that cell *n* belongs to group *t, μ_nt_* is the mean of Poisson distribution followed by the observed read count of cell *n*, which is *y_gn_*. *β*_0_ is an intercept. *β_p_* are coefficients corresponding to covariate *x_np_. δ_t_* is group-specific overexpression. *s_n_* is the size factor^24^ of cell *n*.

Then, the loss function *Q* of each gene is defined using the log likelihood.

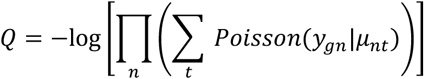

We optimized this loss function using the Adamax optimizer implemented in PyTorch. We used the default learning rate of 2e-3.

As a result, for each gene, we learn how the cells in the datasets are divided into two groups according to the expression modality. Without loss of generality, we assume that the mean expression of group *t*=1 is larger than the mean expression of group *t*=0. We let the indicator variable *I_gn_* ∈ {0,1} denote the cluster assignment of a cell *n* according to gene *g*.

### Sorting out genes of which expression patterns are biologically feasible

To sort out DEGs without taking the group information of the cells as an input, MarcoPolo uses the following multiple crtiteria.

#### (1) Voting system

The voting system prioritizes genes that exhibit a common expression pattern with other genes. Assuming that a gene of interest is truly related to a biological status, it is likely that there are more genes that are co-expressed in a way similar to the gene. In other words, if a gene reflects the structure of the dataset appropriately, its expression pattern will be replicated by other genes several times. We examine how many times the segregation pattern of a gene is repeated by calculating the voting score for each gene as follows.

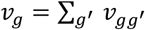

where if 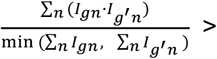 *thres v_gg’_*, = 1, or else *v_gg’_*, = 0 (The default value for *thres* is 0.7)

Thus, the more times a gene is supported by other genes, the higher the gene’s voting score becomes. Note that our formula above calculates what proportion of the cells that express a gene with a smaller on-cell count also expresses a gene with a larger on-cell count. We used this formulation because sometimes, one gene can be a marker gene of a group and another gene can be a marker gene of a subtype of that group. In such a case, we wanted to consider them as supportive of each other in our voting system.

#### (2) Proximity score system

Due to the cell lineage, a hierarchical structure is pervasive in scRNA-seq data. That is, heterogeneity from higher-level grouping determines the global structure, and the cell subtype affects the small signal in the expression. Thus, we assumed that the expression pattern of a gene corresponding to a meaningful biological status tends to align well with the global structure. This idea is similar to the one on which singleCellHaystack is based. For each gene *g,* we calculated the proximity of the on-cells (*I_gn_*=1) in a low dimensional representation space. The underlying intuition is that, if a gene can explain the underlying structure, the cells expressing that gene will be clustered or proximal to each other. We first performed principal component analysis (PCA) of the scaled and normalized data with total 50 components. Then we calculated the proximity score as follows.

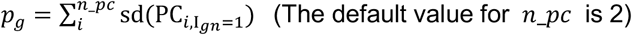

The smaller this score is, we interpret the gene as more informative. Note that although this score tends to capture genes whose on-cells cluster together in the standard clustering approach, our method is not dependent on a specific clustering result. Our method can be interpreted as using the clustering information in PC space in a soft way, so that we can avoid errors induced by fixing the clusters. The way we scale and normalize count data is the same as Seurat. We first divided counts for each cell by the total counts for that cell and multiplied by 10,000. This is then natural-log transformed using log1p. Next, we centered the data to 0 and divided it by the standard deviation. Finally, the values were truncated to 10.

#### (3) Bimodality score system

The bimodality score system prioritizes genes of which expression is bimodal. We measured the discrepancy between the high and low expression components of a given gene using two statistics. First, we compared the log-likelihood of the data under the null hypothesis with a single Poisson distribution and the alternative hypothesis with K=2 Poisson distributions with different means. We have developed a unique statistic defined as the ratio of the two log-likelihoods, namely QQ scores, as follows.

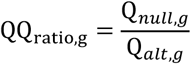

This modeling can look unusual because the subtraction of the two log-likelihoods is more common in other statistical areas. We have found that the ratio statistic fits this problem well because the ratio is independent of the absolute read counts if the on-cell and off-cell count distributions are fixed. Because read counts can differ drastically from gene to gene, we found that this statistic performs well for determining the ranks of multiple genes.

In addition to comparing Q-score, the bimodality score ranking system compares the on-cells’ mean expression value with all cells’ mean expression value. In other words, we compared how much the mean of alternative hypothesis shifts from the mean of the null hypothesis as follows.

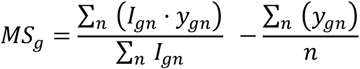

Finally, the bimodality score is obtained by merging the two measures as follows.

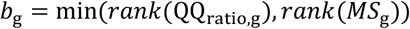

In case two genes have the same rank, the gene with larger fold change is prioritized.

#### (4) Final step of obtaining MarcoPolo score

Finally, we aggregate the abovementioned statistics to select genes with biologically feasible expression patterns. We generate the nonparametric rank-based MarcoPolo score for each gene by combining *v_g_, p_g_*, and *b_g_*.

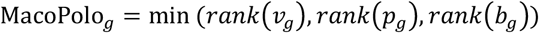

In case two genes have the same rank, the gene with larger fold change is prioritized. After calculating this statistic, we removed outlier genes that satisfy one of the following conditions: (1) log fold change between on-cells and off-cells is < 0.6, or (2) *∑_n_ I_gn_*, the number of on-cells, is < 10 or under the bottom 30th percentile.

### Datasets

#### Embryonic stem cell scRNA data

The Koh et al.^20^ dataset consists of 531 human embryonic stem cells (hESCs) at various stages of differentiation. We extracted the data from the R package DuoClustering2018^25^, which can be installed using Bioconductor package manager. The dataset contains 9 cell types. Among them, we used 8 cell types with both scRNA-seq data and bulk RNA-seq data, which are hESC (day 0), anterior primitive streak (day 1), mid primitive streak (day 1), DLL1+ paraxial mesoderm (day 2), lateral mesoderm (day 2), early somite (day 3), sclerotome (day 6), central dermomyotome (day 5). Koh et al. annotated the cell types through fluorescence-activated cell sorting (FACS).

#### Human liver scRNA data

The MacParland et al.^21^ liver dataset consists of 8,444 cells of 11 cell types collected from 5 patients. We extracted the data from the R package HumanLiver, which can be downloaded from https://github.com/BaderLab/HumanLiver. After clustering cells, MacParland et al. determined the identity of each cluster using known gene expression profiles. We mapped 20 discrete cell populations identified by the authors to 11 unique cell types for our analysis (hepatocytes, ab T cells, macrophages, plasma cells, NK cells, γδ T cells, LSECs, mature B cells, cholangiocytes, erythroid cells, hepatic stellate cells).

#### PBMC 4k scRNA data

We obtained PBMC 4k (peripheral blood mononuclear cell) dataset^22^, namely Zhengmix8eq, from the R package DuoClustering2018. This dataset is a mixture of 3,994 FACS purified PBMC cells of 8 cell types, which are B cells, monocytes, naive cytotoxic cells, regulatory T cells, memory T cells, helper T cells, naive T cells, and natural killer cells.

#### Tabula Muris consortium data

We downloaded Tabula Muris consortium data^23^ from the consortium website (https://tabula-muris.ds.czbiohub.org/). It consists of 20 datasets of different organs and tissues: aorta, bladder, brain (myeloid), brain (non-myeloid), diaphragm, fat, heart, kidney, large intestine, limb muscle, liver, lung, mammary gland, marrow, pancreas, skin, spleen, thymus, tongue, and trachea. They provided datasets obtained by using two kinds of approaches, microfluidic droplet-based 3’-end counting, or FACS-based full length transcript analysis. We used FACS-based datasets which have higher sensitivity and coverage than microfluidic datasets.

#### Simulation data

We generated multiple scRNA-seq simulation datasets using Symsim^19^, a simulator of single-cell RNA-seq experiment. Each dataset contained 1,000 cells with 5,000 genes sequenced. We modulated a parameter, the probability that the gene effect size is not zero, in the simulator. We ran simulations ten times for each combination of parameters with different random seeds. In total, 40 datasets were generated. For the rest of the parameters, we used the tree structure of numerous subpopulations (**Supplementary Figure 6**) using the following setting. The number of extrinsic variability factors (EVFs) was 10. The number of different EVFs between subpopulations (Diff-EVFs) was 5. The mean of a normal distribution from which gene effects are sampled was 1. The parameter bimod, modifying the amount of bimodality in the transcript count distribution, was 1.

#### Data preprocessing

We used only the genes of which mean expression (log-normalized count) value across all cells was in the top 30th percentile. In all the 23 real datasets, meta information of the samples included the t-SNE coordinates. We used them for plotting figures. The size factors of cells were calculated using calculateSumFactors^24^ implemented in scran package^26^.

#### Marker definition

We defined marker genes for each cell type following the same procedure described in Zhang et al^27^. Briefly, for each gene, we sorted the cell types in ascending order based on the mean expression level. We then calculated log fold change between two consecutive types in this order. We then chose the maximum value among the N-1 log fold change values, given N types. After calculating this maximum value for all genes, we used genes with the maximum values in the top 200 as marker genes. Among them, we filtered genes with the maximum log fold change not larger than two-fold for real datasets and four-fold for simulation datasets.

#### AUC calculation

We compared the performance of MarcoPolo with those of HVG methods, singleCellHaystack, and Seurat standard pipelines. For HVG methods (VST or DISP), we used the lists of highly variable genes from Seurat. For singleCellHaystack, we used the default setting of haystack function included in the package. For the standard DEG pipeline with clustering, we used the default parameters of the Seurat package. We used Wilcoxon Rank Sum implemented in FindAllMarkers function to find genes that are differentially expressed.

We sorted the genes by their p-values. We used roc_auc_score function in scikit-learn package to calculate the AUC of each method and dataset pair.

#### Generating MarcoPolo HTML report

To help researchers conveniently interpret and make use of the result for various purposes, we provide the analysis result in the form of an HTML file. For each gene, the output file provides fold change based on the tentative group, a two-dimensional plot of cells, a histogram of expression with annotated group information, and the statistics that were used to prioritize genes. For the histogram, cells with zero expression count were excluded from plotting. In addition, it contains a biological description of each gene that was adopted from the NCBI Gene database^28^. The gene description information was downloaded from NCBI FTP server (https://ftp.ncbi.nih.gov/gene/DATA/GENE_INFO/Mammalia/Homo_sapiens.gene_info.gz).

#### Using the grouping information from MarcoPolo to identify cell types

MarcoPolo provides grouping information according to the bimodality in each gene’s expression as a result. Using this information, we measured how many genes at the top in MarcoPolo gave sufficient information to fully segregate all cell types in each dataset. To this end, we regarded a gene to be the marker of a true cell type label if the gene was expressed in more than 70% of cells of the type. Then we reviewed the genelabel linkage in the order of genes appearing in the MarcoPolo report. When there is a unique gene-label linkage sequence that can be used to discriminate the corresponding label from others, we concluded that the label is identified. For singleCellHaystack, we used the same order in its result. However, as it does not estimate grouping information, we used a hard threshold of 1 for expression value to classify cells into two groups.

#### Using MarcoPolo as a feature selection method to improve the robustness of the clustering process

Since MarcoPolo and singleCellHaystack are designed to pick genes with informative differential expression, we used them as a feature selection method in the standard clustering pipeline. We measured the frequency of successfully clustering populations when MarcoPolo genes or singleCellHaystack genes were employed. We compared three categories of feature selection methods: only using genes selected by standard HVG method, using the mixture of HVG genes and MarcoPolo genes (namely HVG with MarcoPolo), and lastly using the mixture of HVG genes and singleCellHaystack genes (namely HVG with Haystack). That is, for each category of feature selection methods, we repeatedly clustered cells 450 times with different settings (2 HVG methods × 9 HVG numbers × 5 top PC numbers x 5 resolution parameters). We then calculated how many of those 450 trials succeeded in isolating the populations. In each trial, we determined if the two cell types were isolated as follows. We mapped each group of clustering result to a cell type if more than 80% of cells in the group is from the cell type. Then if the proportion of correctly mapped cells of two cell types was larger than 0.8, we concluded that the two cell types were distinguished in the clustering result.

## Declaration

### Ethics approval and consent to participate

Not applicable.

### Consent for publication

Not applicable.

### Availability of data and materials

MarcoPolo is available at the GitHub repository (https://github.com/ch6845/MarcoPolo). It is coded in Python 3.7 using PyTorch v1.4. The required packages are NumPy, SciPy, scikit-learn, and Pandas.

MarcoPolo reports for real datasets used in the analysis are available online.

hESC (Koh et al.) (https://ch6845.github.io/MarcoPolo/Kohinbulk/index.html)

Liver (MacParland et al.) (https://ch6845.github.io/MarcoPolo/HumanLiver/index.html)

PBMC (Zheng et al.) (https://ch6845.github.io/MarcoPolo/Zhengmix8eq/index.html)

The followings are Tabula Muris consortium datasets.

Aorta (https://ch6845.github.io/MarcoPolo/TabulaAorta/index.html)

Bladder (https://ch6845.github.io/MarcoPolo/TabulaBladder/index.html)

Brain Myeloid (https://ch6845.github.io/MarcoPolo/TabulaBrainMyeloid/index.html)

Brain Non-Myeloid (https://ch6845.github.io/MarcoPolo/TabulaBrainNonMyeloid/index.html)

Diaphragm (https://ch6845.github.io/MarcoPolo/TabulaDiaphragm/index.html)

Fat (https://ch6845.github.io/MarcoPolo/TabulaFat/index.html)

Heart (https://ch6845.github.io/MarcoPolo/TabulaHeart/index.html)

Kidney (https://ch6845.github.io/MarcoPolo/TabulaKidney/index.html)

Large intestine (https://ch6845.github.io/MarcoPolo/TabulaLargeIntestine/index.html)

Limb muscle (https://ch6845.github.io/MarcoPolo/TabulaLimbMuscle/index.html)

Liver (https://ch6845.github.io/MarcoPolo/TabulaLiver/index.html)

Lung (https://ch6845.github.io/MarcoPolo/TabulaLung/index.html)

Mammary Gland (https://ch6845.github.io/MarcoPolo/TabulaMammaryGland/index.html)

Ma rrow (https://ch6845.github.io/MarcoPolo/TabulaMarrow/index.html)

Pancreas (https://ch6845.github.io/MarcoPolo/TabulaPancreas/index.html)

Skin (https://ch6845.github.io/MarcoPolo/TabulaSkin/index.html)

Spleen (https://ch6845.github.io/MarcoPolo/TabulaSpleen/index.html)

Thymus (https://ch6845.github.io/MarcoPolo/TabulaThymus/index.html)

Tongue (https://ch6845.github.io/MarcoPolo/TabulaTongue/index.html)

Trachea (https://ch6845.github.io/MarcoPolo/TabulaTrachea/index.html)

### Competing interests

Buhm Han is CTO of Genealogy Inc.

### Funding

This work was supported by the National Research Foundation of Korea (NRF) (Grant number 2019R1A2C2002608) funded by the Korean government, Ministry of Science, and ICT. BH and KJ were supported by the Creative-Pioneering Researchers Program funded by Seoul National University (SNU).

---

**Supplementary Figure 1.**
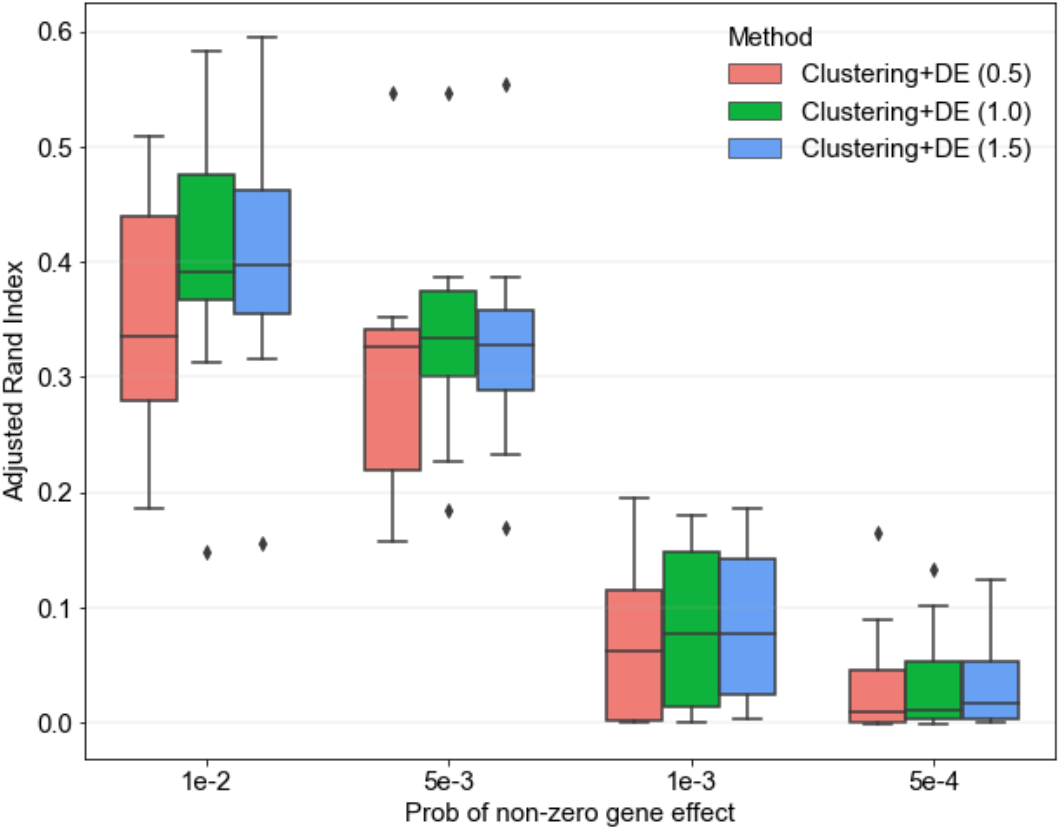
Adjusted rand index of simulation datasets when changing probability of nonzero type-specific expression effect sizes. The number inside parenthesis denotes the resolution parameter of the clustering algorithm

**Supplementary Figure 2.**
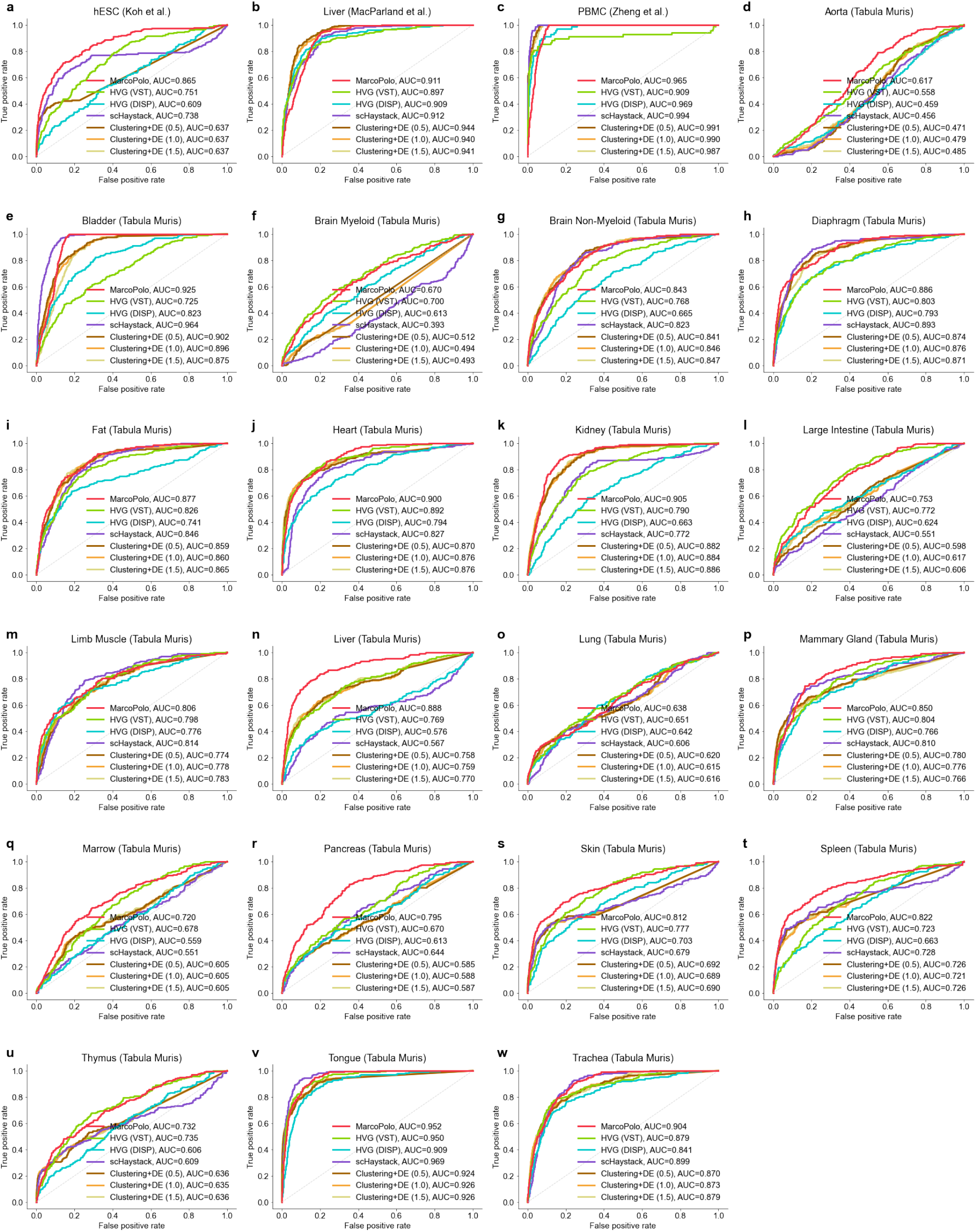
ROC curves and their AUCs of gene lists generated using different methods. The gene lists were obtained by using MarcoPolo, HVG methods, singleCellHaystack, or standard DEG pipeline with clustering. For standard DEG pipeline, the number inside parenthesis denotes the resolution parameter of the clustering algorithm

**Supplementary Figure 3.**
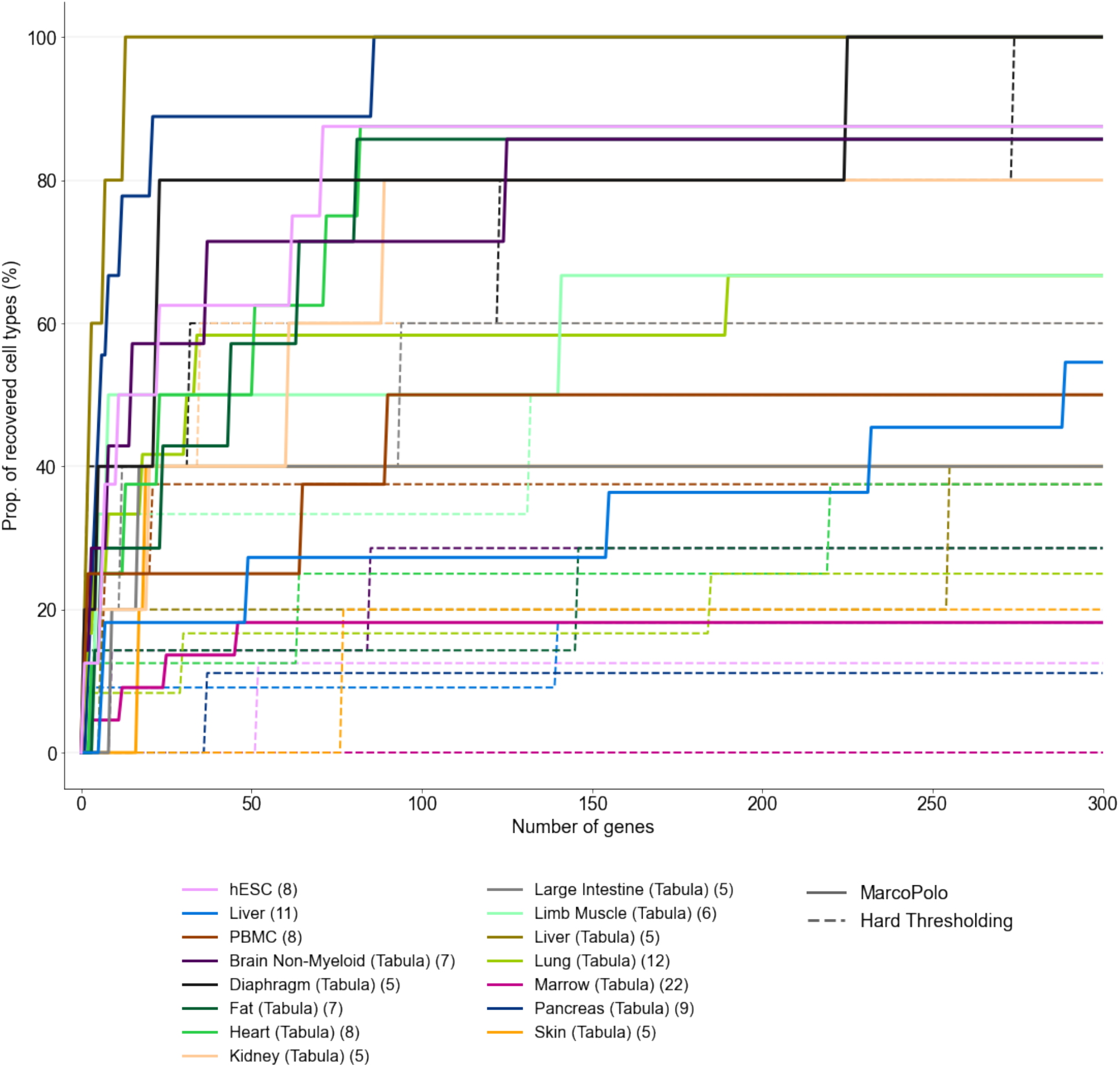
The proportion of recovered cell types in the datasets by using MarcoPolo’s grouping information. The grouping information of each gene was reviewed according to its rank assigned by MarcoPolo. Only datasets containing more than 4 cell types are shown. The number inside the parenthesis beside the dataset name denotes the number of cell types in each dataset.

**Supplementary Figure 4.**
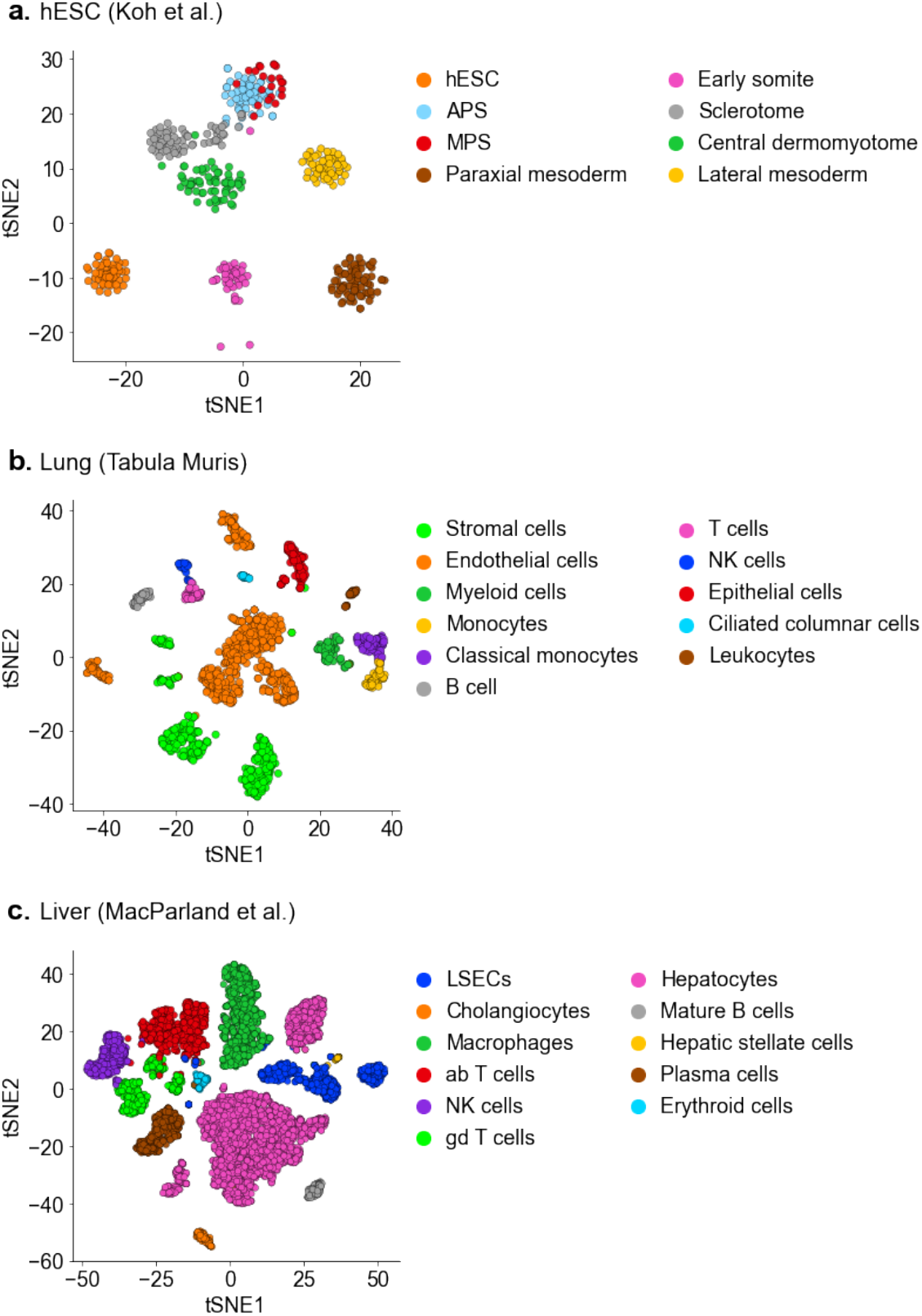
t-SNE plots included in the downloaded datasets. **(a)** hESC dataset (Koh et al.) The hESC dataset was preprocessed by Duò et al^25^. **(b)** the lung dataset (Tabula Muris consortium). **(c)** the liver dataset (MacParland et al.)

**Supplementary Figure 5.**
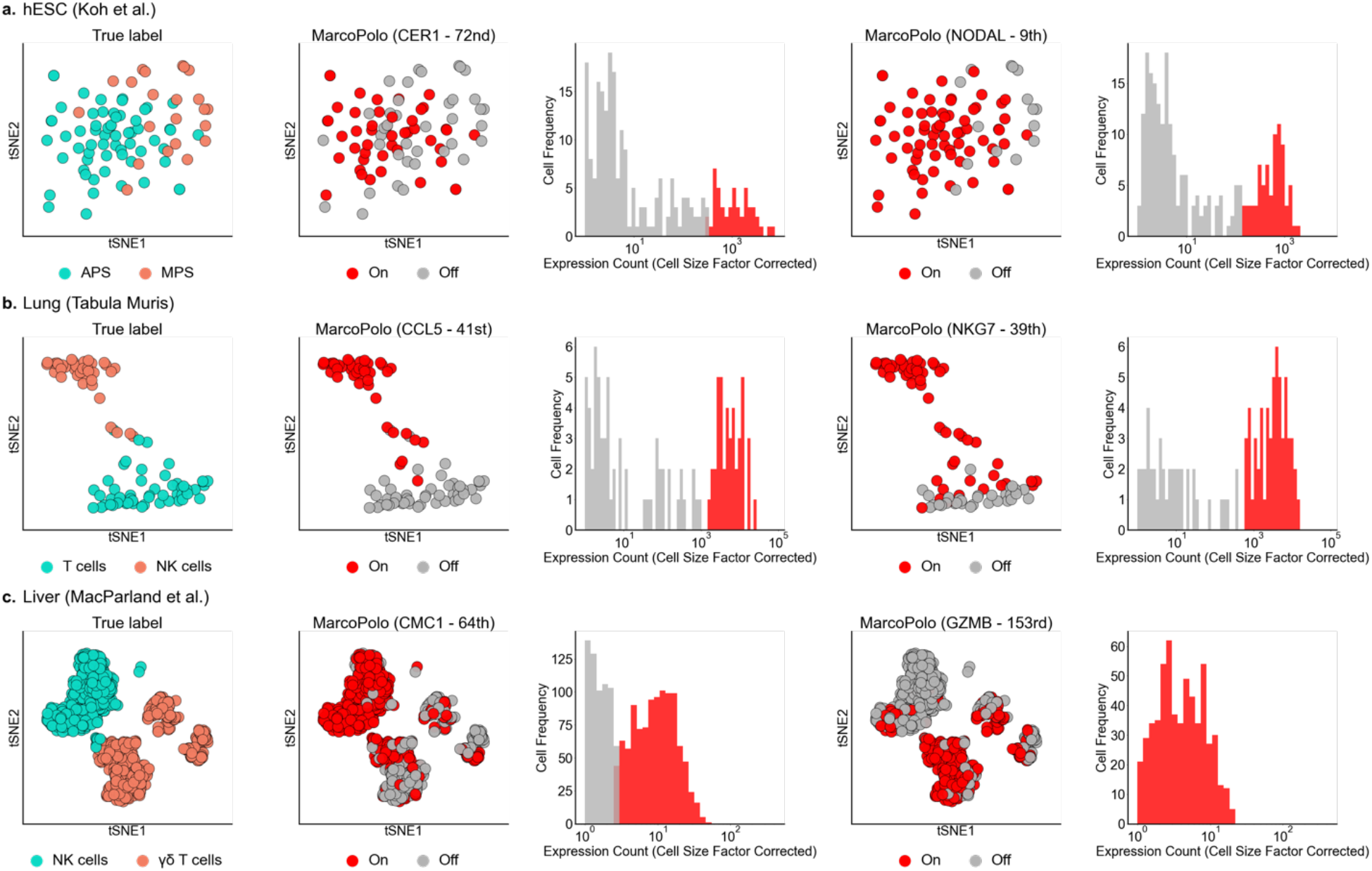
Cell types distinguished by MarcoPolo for genes in Table1. They are cell types that were not distinguished using the default option of the standard clustering pipeline. Exceptionally, we used the clustering resolution parameter of 2.0 (default: 1.0) to obtain a finer clustering result. We plotted cells on the precalculated t-SNE coordinates that were included in the downloaded datasets. For each dataset, we only showed cell types of interest. In the t-SNE plot in the first column, we colored cells by their true cell-type label. In the next t-SNE plot, we colored cells based on MarcoPolo grouping. For each gene, MarcoPolo learns how to divide cells in the given dataset into two groups according to their expression modality. The cells belonging to the on-cell group (i.e., the group of higher gene expression) were colored red. In the histogram, the expression pattern of each gene was shown. Cells with zero expression count were excluded from plotting. The expression count was divided by cell-specific scaling factors (also known as size factors). **(a)** the hESC dataset of Koh et al. **(b)** the lung dataset of Tabula Muris consortium **(c)** the liver dataset of MacParland et al.

**Supplementary Figure 6.**
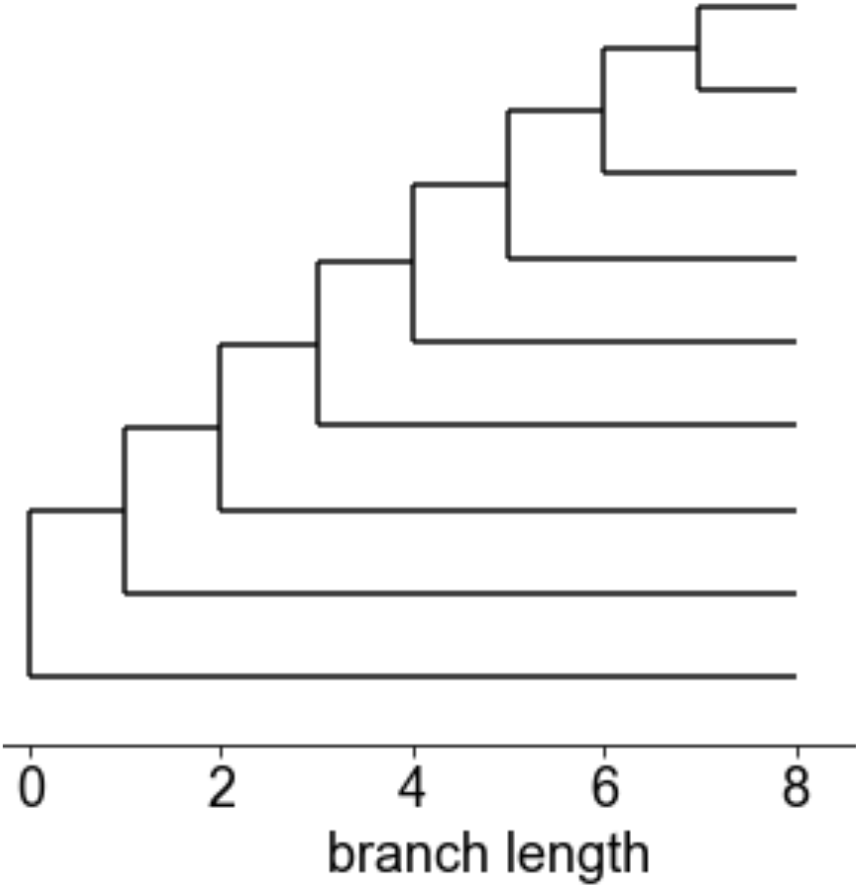
Structure of cell subpopulations used to generate simulation datasets.

## Notes

https://github.com/ch6845/MarcoPolo

### Competing Interest Statement

Buhm Han is the CTO of Genealogy Inc.

